# Short-term antibiotics, induced magnetic field, and magnetic sensitivity in a migratory songbird

**DOI:** 10.1101/2024.09.19.613835

**Authors:** Gaya Sherf, Anat Levi, Eviatar Natan, Yoni Vortman

**Affiliations:** Hula Research Center, Department of Biotechnology, Tel-Hai College, Israel; Hula Research Center, Department of Animal Sciences, Tel-Hai College, Israel; The Aleph Lab, Oxford, UK; MIGAL – Galilee Research Center, Israel

**Author notes:** Corresponding author: Gaya Sherf.

**Keywords:** Magnetotactic bacteria (MTB), Magnetic sensing, Passerines, Helmholtz coils, Symbiotic magnetic sensing

## Abstract

The use of magnetoreception is widespread in the animal kingdom. Although much effort has been invested in unraveling the receptor and the mechanism behind this ability, the results remain inconclusive. The recently proposed ’symbiotic magnetic sensing hypothesis’ suggests that magnetotactic bacteria symbiotically act as ‘magnetic sensors’ for the host. In this study, we examined the changes in magnetic orientation of Eurasian reed warblers (*Acrocephalus scirpaceus*) placed in Emlen funnels under the influence of antibiotics, in response to a manipulated magnetic field. We found that reed warblers of the control group showed no significant magnetic orientation when placed in a manipulated magnetic field. In contrast, reed warblers placed in an unplugged coil showed significant magnetic southwest orientation. Furthermore, reed warblers exposed to short-term antibiotic treatment (12 h) demonstrated significant magnetic orientation (southwest, regardless of season) and significantly lower directional variance. We hypothesize that individuals from the control group identified the electromagnetic noise induced by the artificial magnetic field. Taken together, this unexpected result may demonstrate the sensitivity of passerines to electromagnetic noise and a possible damping effect of antibiotics on this sensitivity. Assuming our interpretations are correct, we consider this result as indirect experimental support for the symbiotic magnetic sensing hypothesis.

## Introduction

Many organisms, including invertebrates, vertebrates, and bacteria (Wiltschko & Wiltschko, 2012), have demonstrated an ability to use the earth’s magnetic field for orientation and navigation. Most organisms use the inclination angle of the earth’s magnetic field (Putman et al., 2011; Wiltschko & Wiltschko, 1972; Wynn et al., 2022)—the angle formed where the magnetic field lines intersect the surface of the earth— which ranges from 0° at the equator to 90° at the poles. Since it varies with latitude, it can be used as an indication of location for migratory species (Lohmann & Lohmann, 1994; Wynn et al., 2022). Although much effort has been made for over 50 years to discover the identity of the receptor and the mechanism of magnetoreception in animals, they still remain unknown (Nordmann et al., 2017).

A major challenge in revealing the identity of the receptor and mechanism is a methodological difficulty that arises from the animal’s ability to ignore the magnetic sense. Although the earth’s magnetic field is omnipresent, the use of it as a navigational cue is usually at the bottom of the compasses hierarchy (Gould, 1998; Johnsen et al., 2020). Furthermore, there seems to be great variation in species’ sensitivity to the magnetic field. Several migratory bird species have exhibited sensitivity to oscillating magnetic fields, where some species are more sensitive than others. Moreover, the birds express this sensitivity as a lack of orientation, demonstrating a noticeable effect that is behaviorally a “negative” result (Bojarinova et al., 2023). Another challenge in the research of magnetoreception is that as humans, we lack the perception of the earth’s magnetic field (but see Wang et al. 2019). Thus, designing an experiment would be an unintuitive and very complex task (Nordmann et al., 2017).

Taken together, with respect to many experiments in the field of magnetoreception, the tested animal’s behavior and the subsequent results might be weak and demonstrate low statistical power. The tested individuals will select a wide range of headings, in turn leading to an unreliable group mean azimuth and increase in directional variance (Johnsen et al., 2020), or even unreproducible experiments (Winklhofer et al., 2023).

In light of the above, the magnetic sense remains the only sense without a known sensor; while various hypotheses have been proposed regarding the underlying mechanism (Hore & Mouritsen, 2016; Kirschvink et al., 2001; Natan & Vortman, 2017; Nordmann et al., 2017) all currently remain hypotheses. Recently, the symbiotic magnetic sensing hypothesis has been proposed, suggesting that magnetotactic bacteria (MTB) symbiotically act as ‘magnetic sensors’ for the host (Natan & Vortman, 2017). As opposed to other organisms that use the earth’s magnetic field for orientation and navigation, the only proven magnetoreception mechanism is that of MTB (Goswami et al., 2022) or based on MTB (Monteil et al., 2019).

The MTB are aquatic, gram-negative prokaryotes, diverse in habitat, morphology, and taxonomy. These bacteria mineralize ferromagnetic crystals inside an organelle called a magnetosome and use it to align themselves along the earth’s magnetic field lines (Lefevre & Bazylinski, 2013).

In recent years, some supporting evidence for the symbiotic magnetic sensing hypothesis has accumulated. A symbiosis has been demonstrated between a eukaryotic marine protist and MTB, in which the protist gains the magnetic moment from its resident MTB (Monteil et al., 2019). Furthermore, a database of metagenomic surveys, called MGRAST (Meyer et al., 2008), was examined, focusing on samples that were derived from organism-associated environments, to learn about the prevalence of MTB in animal tissues. The presence of MTB was observed in more than 4000 different samples taken from various animal species. Samples taken from the loggerhead sea turtle (*Caretta caretta*) and various penguin species contained similar MTB, mostly *Candidatus Magnetobacterium bavaricum*. Additionally, in samples taken from mammalian species, brown bats (*Myotis*) and Atlantic right whales (*Eubalaena glacialis*) had similar MTB, mostly *Magnetospirillum* and *Magnetococcus*. These findings allow us to assume that MTB are abundant in the microbiome of many animal species, and that phylogenetically related species from diverse habitats may present a relatively similar composition of MTB. Moreover, this widespread presence of MTB among animals makes the symbiotic magnetic sensing hypothesis more probable than first thought (Natan et al., 2020).

A recent study showed that antibiotic treatment caused a lack of orientation in a migratory species (Eurasian reed warbler (*Acrocephalus scirpaceus*)) in an Emlen funnel experiment (Werber et al., 2022). However, in this experiment the funnels were not placed in an artificial magnetic field, and while all celestial cues were unavailable to the birds, this experiment did not specifically isolate the magnetic sense. Furthermore, a surprising result of Werber et al. (2022) was a significant southwest spring orientation of the control group of birds.

Here, following the results of Werber et al. (2022) we conducted a migratory restlessness experiment using an artificially induced magnetic field in Helmholtz coils (Packmor et al., 2021; Wiltschko & Wiltschko, 1972). We examined the effect of very short antibiotic treatment on the migratory restlessness orientation of reed warblers held in a manipulated magnetic field. We analyze the results in light of previous research and discuss the effect of short-term antibiotic treatment on magnetic sensitivity with respect to the symbiotic magnetic sensing hypothesis.

## Materials and Methods

### Study site and experimental periods

The study was conducted at the Hula Research Center, Hula Valley, Israel (33°06’43.1’’N 35°35’07.7’’E) during two spring migration seasons (March–May of 2021 and 2022) and one autumn migration season (September–October of 2021). The Hula Valley, located in the middle of the Eurasian African flyway, serves as a prominent stopover site for migratory birds, making it an ideal research area for studying migratory avian species and migration dynamics.

### Study animal

The Eurasian reed warbler (*Acrocephalus scirpaceus* (Hermann, 1804)) was selected as the study species. This species is a nocturnal migratory passerine, undertaking seasonal migrations from Europe and west Asia to Africa during autumn and returning to breeding grounds during spring. The breeding range of this species includes Israel (Pearson et al., 2002). The high abundance of Eurasian reed warblers passing through the Hula Valley, as well as their extensive use as a model species in previous studies on magnetic compass orientation and navigation (Chernetsov et al., 2017; Packmor et al., 2021; Werber et al., 2022), led us to choose this species for the study.

### Experimental setup

For the initial two seasons, the experiment was conducted in a small building made partly of metal. Therefore, adjustments were made using our Helmholtz coils to overcome the potential changes to the magnetic field, as will be described later. In the subsequent seasons we moved the experimental setup to a wooden cabin, which had no effect on the magnetic field inside the room or inside the coils. Magnetic field measurements were taken in the position of the birds with a Lake Shore Model 460 Gaussmeter.

The Helmholtz coils allow artificial manipulation of local magnetic fields, e.g., inversion of magnetic north to magnetic south or west (Wiltschko and Wiltschko 1972; Emlen et al., 1976). This well-established experimental protocol is to date the most prominent methodological paradigm (Packmor et al., 2021). Four Helmholtz coils were custom-made by Kanfit Ltd and fed with power suppliers (Nice Power R-SPS605 60V 5A regulated DC Power Supply and APS3005S-3D regulated DC power supply) to create four different local magnetic fields: one with magnetic north inverted to magnetic south, two with magnetic north inverted to magnetic west, and one with no inversion (but still turned on), to resemble ‘true’ north. Four Emlen funnels (Emlen & Emlen, 1966) were placed within each coil, allowing a maximum of 16 individual birds to be tested in each experimental trial.

During the experiments conducted in the metal building, the net magnetic field was corrected by manipulating the intensity of the magnetic field in the horizontal axis, thereby achieving the appropriate net induced inclination angle (local inclination angle ∼49°, angles produced ranged from 38° to 57°), even with a slight compromise in magnetic field intensity (∼41–52 µT). Upon relocating the experimental setup to the designated wooden cabin, such compromises in both inclination angle (local inclination angle ∼49°, angles produced ranged from 48° to 51°) and magnetic field intensity (45–50 µT) were not necessary. Furthermore, during spring 2022, the fourth coil (north coil) was left unplugged to simulate the natural magnetic field (there were no detectable anomalies in the wooden housing).

Considering the consistent findings across all three experimental seasons, we hypothesized that the magnetic field exhibited instability, resulting in high noise levels. To investigate the presence of electromagnetic noise within the coils during operation, we recorded the magnetic field using a TFM1186 3-axis Fluxgate Magnetometer (Metrolab). For each coil, two one-minute measurements were captured at a frequency of 100 Hz: one with the coil activated and the other with the coil deactivated.

### Lighting conditions

In the first season, experiments were conducted solely with natural lighting from outside the building, resulting in variation in light levels within the room each night. To address this, a controlled artificial lighting setup was introduced due to the low expression of migratory restlessness among the birds.

During the day, the birds were exposed to natural daylight entering the room through the windows. At the time of the experiment the curtains were closed and four HiraLite LED light bulbs were turned on. The HiraLite LED light bulbs emit a broad spectrum of light that closely resembles natural light.

Each coil had one lightbulb installed above it, facing the ceiling, to create an even distribution of the light around the room and prevent the birds from flying towards a certain light source. The lightbulbs were connected to a dimmer and covered with a net to maintain the light intensity at 0.04 lux (2.5±0.25 mW m^−2^) (Schwarze et al., 2016; Zapka et al., 2009). Light measurements were made at the position of the birds with an ILT2400 hand-held light meter/ optometer.

### Experimental procedure

#### Bird capture

Eurasian reed warblers were captured early in the morning at the Hula ringing station using mist nets. In the autumn of 2021, an additional capturing site was established in a reed area located 2.3 km north of the ringing station. Birds selected for the experiment had a fat score of 3 or higher, indicating their readiness to migrate soon (Werber et al., 2022). During spring migration, a wing length threshold exceeding 66 mm was also implemented to avoid selecting birds from the local population that had already reached their breeding grounds (Pearson et al., 2002). Suitable individuals were divided into Control and Treatment groups and placed inside cloth bags for transportation to the research center.

#### Treatment administration

Upon arrival at the research center, each reed warbler was administered either 4.6 µl water (Control group) or 4.6 µl Enrofloxacin (50 mg/ml solution), also known as Baytril (Treatment group). Enrofloxacin was chosen based on a previous laboratory experiment where it demonstrated lethality against various types of MTB (Werber et al. 2022).

Unlike the previous experiment, in which birds were kept for three days (Werber et al. 2022), the duration of captivity for the current experiments was reduced to 12–16 h from capture to release to increase sample size and minimize stress. To increase the probability that the antibiotics will have an effect in such a short time, we increased the usual dose for mean reed warbler body weight by 15% in comparison to our previous research (Werber et al. 2022). Upon arrival at the research center, each reed warbler was orally administered the first dose and was placed in a metal bird cage (30×23×40 cm) with access to water. After two seasons, the metal cages were replaced with wooden cages of the same size to minimize potential magnetic interruptions. The second dose was administered 5 h after the initial dose. At any stage of the day, a bird that seemed weak or stressed was released.

### Experimental setup and recording

Preparations for the experiment started at sunset. Helmholtz coils were turned on with the appropriate current and each reed warbler was placed inside an Emlen funnel which was then covered with a dark glass. (Note: in the first two seasons, Emlen funnels located in parts of the coil with extreme inclination values of 38–44° or 53–57°, were excluded).

The experiment started 40 min after sunset. Birds were filmed for 80 min: 20 min for the birds’ adjustment and 60 min for recording. We used one HIKvision 2.8 IR camera per coil. The cameras were connected to an HIKvision HD DVR hard drive. At the end of the experiment the birds were released.

### Video data analysis

For the video analysis we used DeepLabCut (version 2.0) (Mathis et al., 2018; Nath et al., 2019). We labeled our point of interest, i.e., the tip of the bird’s beak, in 810 frames taken from 57 videos; 95% of these frames were used for algorithm training. We used a ResNet-50 based neural network with default parameters and trained it for 400,000 iterations. To validate network performance, we conducted one shuffle and found a test error of 2.18 pixels and a train error of 1.74 pixels (image size was approximately 360×340 pixels). We then applied a p-cutoff of 0.6 to condition the X- and Y-coordinates for subsequent analysis. Using this trained network, we analyzed the videos of 83 individuals that exhibited jumping behavior inside the funnels. The resulting coordinate data, in CSV format, was analyzed to determine the azimuth of each jump and calculate the mean azimuth for each individual relative to the induced north, using R 4.1.2 (R Core Team, 2021) and ‘circular’ and ‘CircStats’ packages.

Initially, we needed to define a jump as the exit of the bird’s beak (specific X,Y location) beyond a circle. The circle was defined as halfway between the inner circle (depicted in white) where the bird rests and the outer circle (the edges of the Emlen funnel). This was conducted to create a threshold of the bird’s beak coordinates to ensure a true read of jump, while excluding false counts due to image distortion. Simultaneously to counting the number of jumps, the induced north was inserted as a vector from the center of the circle to a specific point represented by the end of the wooden stick (see Experimental setup and recording). Ultimately, the angles of each of the jumps were calculated and corrected relative to the induced north. This process was performed for all individuals in both Control and Treatment groups. The mean azimuth for each experimental group was then calculated.

For validation purposes, for each individual the direction and time of a number of 5–7 random jumps from the original video were compared to the automated computer analysis. Due to high occurrence of individuals jumping backwards in spring 2022 we decided to manually analyze the videos of this specific season, using a transparent azimuth meter (“army protractor”). We compared the automated DeepLabCut and manual analysis, for individuals whose data was analyzed by both methods (17 out of 27, spring 2022). Four individuals that had a high number of reverse jumps, exhibited a difference between the two azimuth values of >90°, while the rest had a an almost identical fit between the values of the automated analysis and the manual analysis (see Figure S1 in the supplementary information).

### Statistical analysis

We used Rayleigh’s Z test to examine whether individuals and groups exhibited significant directionality. To compare the variance in directional tendencies between Control and Treatment groups we used Levene’s test for circular data (Landler et al., 2021). Finally, we used the χ² test and t-test to examine the difference in individual numbers of jumps between groups. Statistical analysis was conducted using R 4.1.2 ‘circular’ and ‘CircStats’ packages (R Core Team 2021) and JMP V15 (SAS institute).

### Ethical Note

We chose the reed warbler as our model species as this is the most abundant migrant caught in the Hula valley ringing station. This had benefits both from a conservation and an animal welfare perspective. Conservation wise, we experimented with a negligible fraction of the population. From an animal welfare point of view this created two major benefits: first, from the captured birds, we choose only individuals with high fat load (see methods above). This enabled the birds to cope with the stress imposed by the experiment. Second, the fact that enlarging the sample size is relatively simple, we released immediately all individuals which showed even slight signs of stress during captivity (e.g. repeated movements or atypical postures). Last, we created a protocol with a minimum captivity time (of about 12 hours). The 12 hour food deprivation is a noon issue, as birds with high fat load which are preparing to initiate migration many times are at a fasting stage (gut reduction), and are able to migrate several consecutive days without food (McWilliams and Karasov 2005). At the end of the experiment all individuals were released immediately back to their natural habitat, the Hula Valley. All research was done under animal welfare assurance and permits from the Israel Nature and Park authority (permit number: 42737) and the Israel national animal Care and Use Committee (IACUC permit number: 0005357).

## Results

Over all three seasons, out of the 342 reed warblers tested in Emlen funnels inside an induced magnetic field, 73 exhibited migratory restlessness inside the funnels while 61 exhibited significant within-individual directionality.

Similar to previous experiments in our lab (Werber et al. 2022), there was no significant effect of antibiotics on activity levels in spring 2021 and autumn 2021. Moreover, there was no significant difference in the proportion of individuals who expressed migratory restlessness or the number of jumps among individuals who expressed migratory restlessness (Table 1). However, in the last season (spring 2022) there was a significant effect of antibiotics on both the proportion of individuals who expressed migratory restlessness and the number of jumps within the Emlen funnel (Table 1).

**Table 1.** Proportion of individuals who expressed migratory restlessness and the number of jumps by these individuals.

| Season | Treatment | Proportion |  |  | Number of jumps |  |  |  |  |
| --- | --- | --- | --- | --- | --- | --- | --- | --- | --- |
| | | n/N | $\chi^2$ | P | N | Mean | SE | t | P |
| Spring<br>2021 | Control | 13/98 | 0 | 1 | 13 | 14.9 | 3.8 | 0.46 | 0.65 |
|  | Antibiotics | 13/98 |  |  | 13 | 17 | 2.6 |  |  |
| Autumn<br>2021 | Control | 11/72 | 0.25 | 0.61 | 11 | 38.3 | 9.2 | -1.35 | 0.19 |
|  | Antibiotics | 13/72 |  |  | 13 | 24.3 | 4.5 |  |  |
| Spring<br>2022 | Control | 16/88 | 4.05 | 0.04 | 16 | 70.4 | 17.3 | -3.56 | 0.002 |
|  | Antibiotics | 7/88 |  |  | 7 | 7.8 | 2.4 |  |  |

In the first season, both Control and Treatment groups showed no significant directionality, but the Treatment group demonstrated a non-significant trend towards southwest (spring 2021: Control (n=9, r=0.321, p=0.4), Treatment (n=11, r=0.405, p=0.16); Figure 1a and b, respectively). In autumn 2021 and spring 2022, the Control group again showed no significant directionality (autumn 2021: Control (n=11, r=0.185, p=0.69), spring 2022: Control (n=15, r=0.096, p=0.874); Figure 1c and e, respectively). Birds from the Treatment group again showed marginally significant directional orientation towards southwest (autumn 2021: Treatment (n=11, r=0.461, p>0.094); Figure 1d, and spring 2022: Treatment (n=4, r=0.831, p=0.052); Figure 1f).

**Figure 1:**
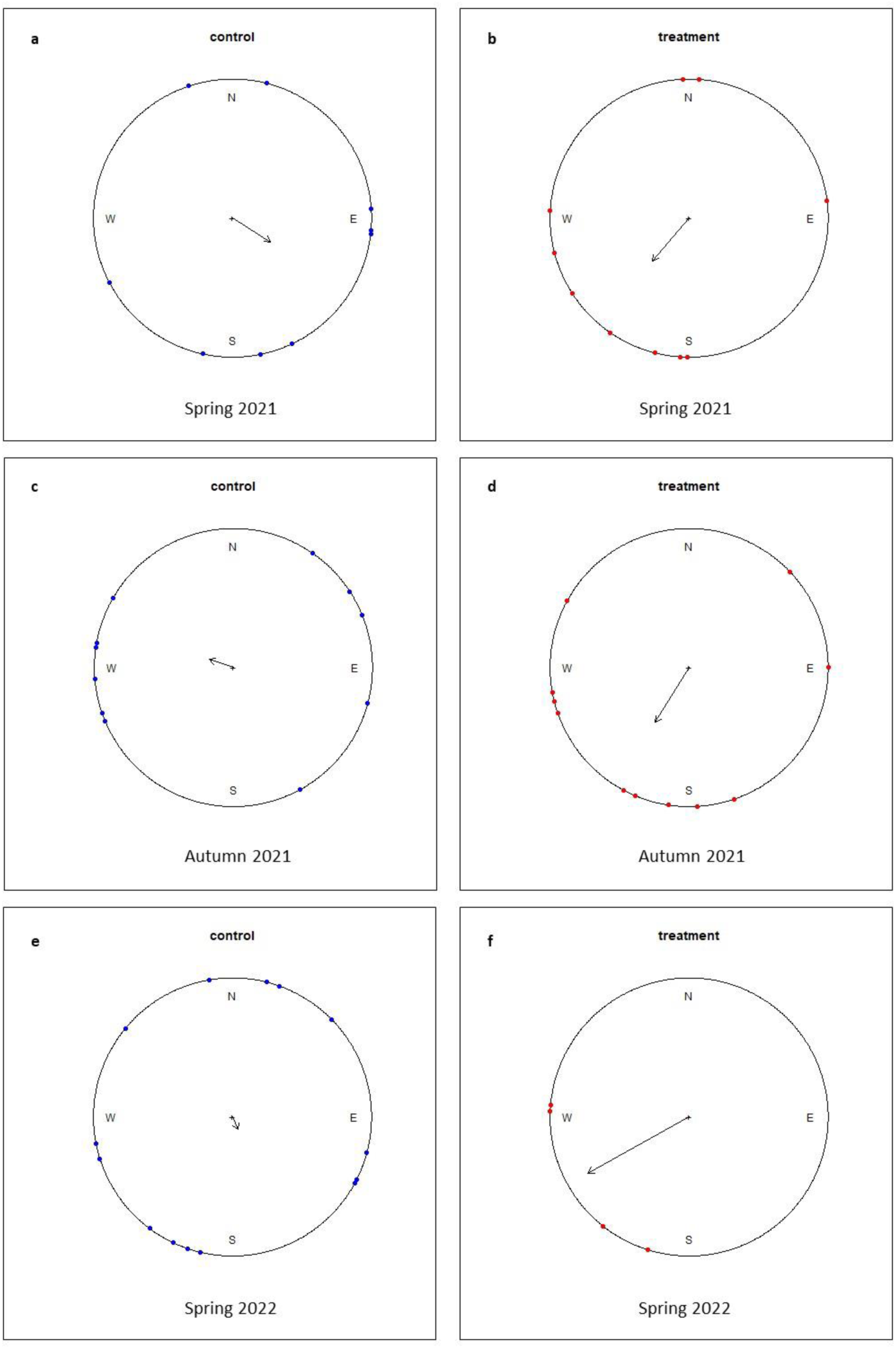
Orientation of treated (red dots) and untreated (blue dots) reed warblers in all three experimental seasons: spring 2021 (a and b), autumn 2021 (c and d), and spring 2022 (e and f). Each dot on the circle represents the average jumping azimuth of an individual inside an Emlen funnel. The direction of the arrow represents the mean orientation of the group and the length of the arrow represents the Rayleigh test R value (ranges from 0 to 1).

The lack of orientation in the Control group on one hand, and the southwest trend of the short-term antibiotic Treatment group on the other hand, repeated itself in all three seasons. This puzzling result could be explained by sensitivity of the individuals in the Control group to electromagnetic noise. Accordingly, we measured the electromagnetic noise within the Helmholtz coils (i.e. the fluctuations in the magnetic field over time). When sampling the magnetic field at a frequency of 100 Hz, we found a significant difference in the variance of the magnetic field between the induced magnetic field and the ambient magnetic field (when the power supply/coil was off), Levene’s test: F_1,1240_ = 35049.74, P < 0.0001. The standard deviation of the ambient magnetic field intensity was 0.004, while the induced magnetic field in the Helmholtz coils had a standard deviation of 0.01 (more than twice the magnitude of the electromagnetic noise, see real time SD graph, Figure S2, supplementary material).

In line with the above we reanalyzed our data with respect to whether the Helmholtz coils were turned on or off. Furthermore, since the Treatment group showed a repeated trend of marginally significant southwest orientation, we reanalyzed the data in two ways: 1) merging all three experimental seasons and 2) merging the two springs, examining the effects of both antibiotics and the induced electromagnetic noise on orientation.

Separately analyzing the unplugged coil created a very limited sample size (4–6 individuals). However, compared to the lack of orientation of the Control group in all three seasons (Figure 1a, c, and e), individuals examined in the unplugged coil from spring 2022 showed significant southwest directionality, showing high R values when analyzing all 6 individuals (Figure 2) or only individuals from the Control group (Figure 2, blue dots), (Control and Treatment: n=6, r=0.76, p=0.02; Control: n=4, r=0.71, p=0.12).

**Figure 2:**
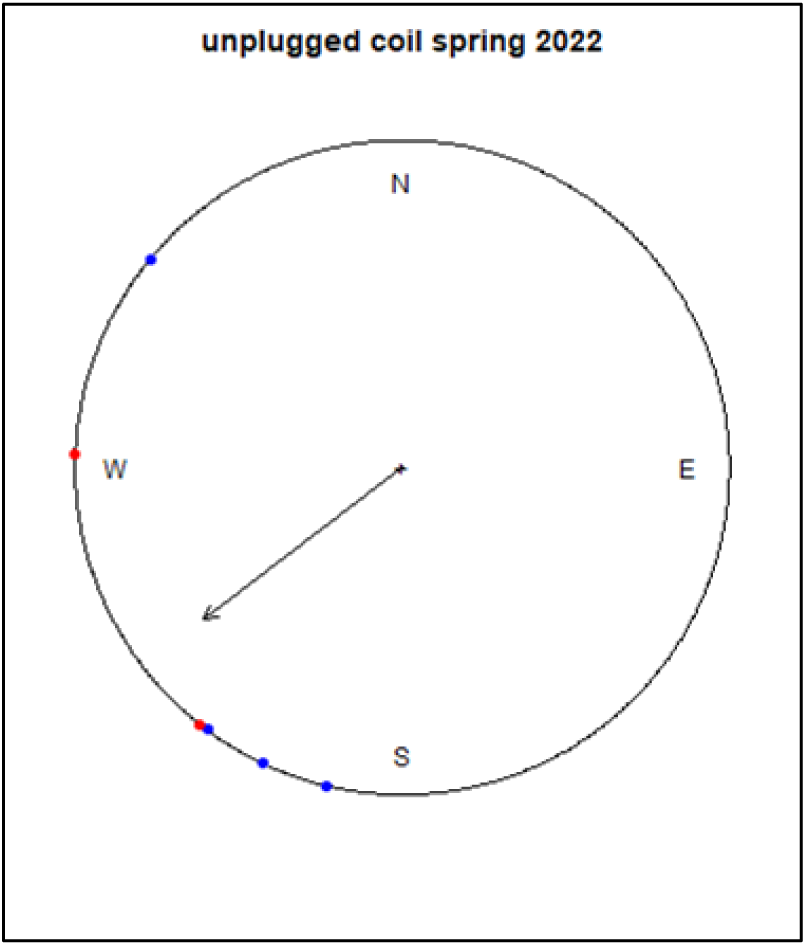
Orientation of treated (red dots) and untreated (blue dots) reed warblers inside unplugged (off) coils (spring 2022). Each dot on the circle represents the average jumping azimuth of an individual inside an Emlen funnel. The direction of the arrow represents the mean orientation of all individuals and the length of the arrow represents the Rayleigh test R value (ranges from 0 to 1).

When merging all three seasons together, but analyzing only individuals placed in active coils, the Control group again demonstrated non-significant directionality (n=31, r=0.1, p=0.735; Figure 3a), while the Treatment group showed significant southwest directionality (n=24, r=0.457, p=0.005; Figure 3b). This result repeated itself when analyzing only the two merged springs: non-significant directionality in the Control group (n=20, r=0.255, p=0.275, Figure 3c) and marginally significant directional orientation in the Treatment group (n=13, r=0.46, p=0.061, Figure 3d).

**Figure 3:**
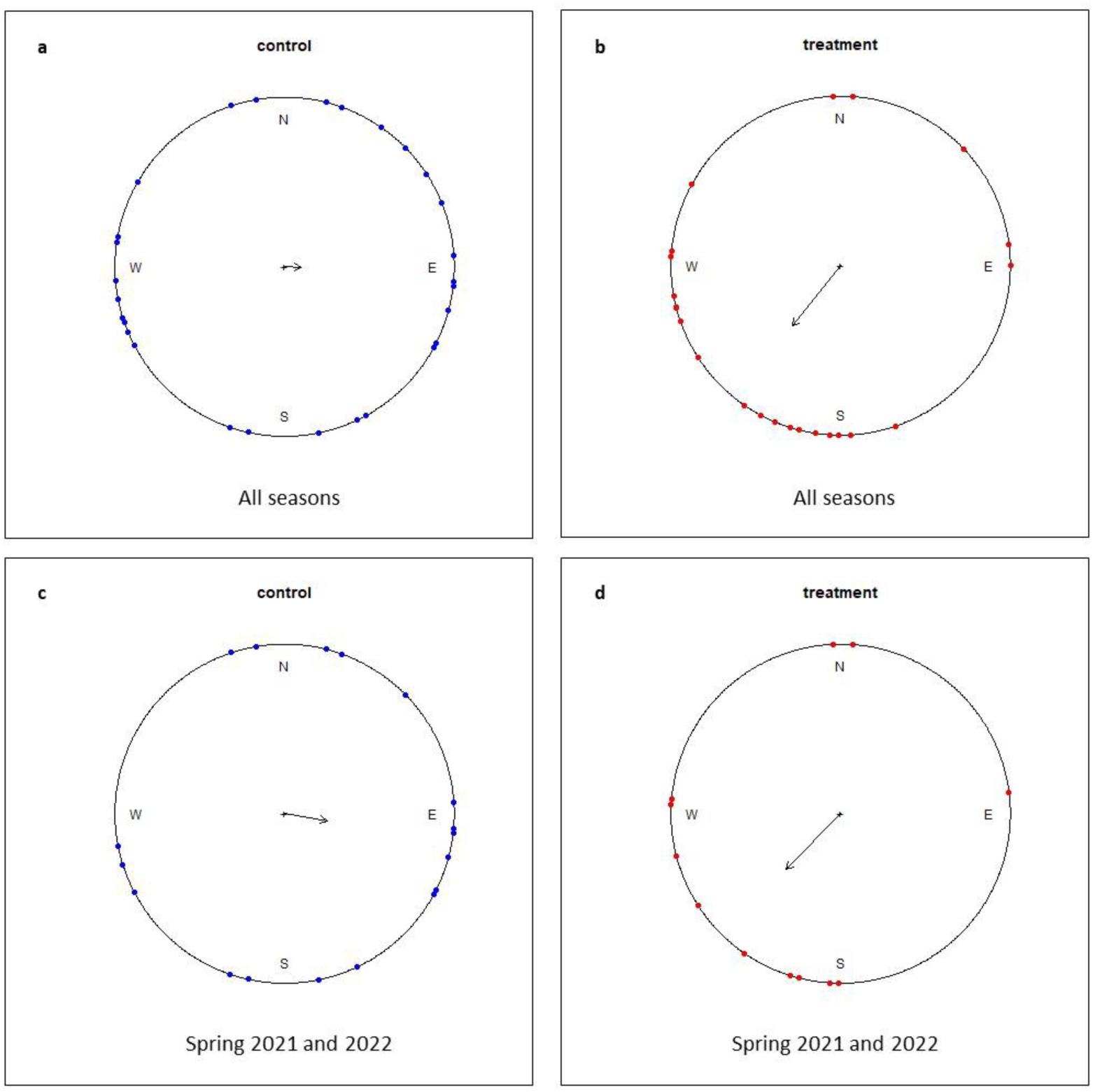
Orientation of treated (red dots) and untreated (blue dots) reed warblers of all three seasons merged (a and b), and two spring seasons merged (c and d). In this analysies we included only individuals examined in a manipulated magnetic field. Each dot on the circle represents the average jumping azimuth of an individual inside an Emlen funnel. The direction of the arrow respresents the mean orientation of the group and the length of the arrow represents the Rayleigh test R value (ranges from 0 to 1).

## Discussion

As mentioned above, the combined use of orientation cages inside an electromagnetic coil system is to date the most prominent methodological paradigm for studies on magnetic compass orientation and navigation, and has been used in many previous studies (Chernetsov et al., 2017; Packmor et al., 2021; Wiltschko & Wiltschko, 1972; Winklhofer et al., 2023). Yet, the results in all of our three experimental seasons repeated themselves, showing no significant directionality in the Control group and marginally significant directional orientation towards the southwest in the Treatment group. The variance in directional tendencies in the Control group was larger, once again indicating a lack of significant group orientation.

Despite the lack of significant directionality of the Control group, in each season separately almost all active individuals showed significant within-individual directionality (regardless of treatment). This means that each individual was consistent in its direction of jumping, perhaps implying that a decision was made by the individual, but the group analysis suggests that the magnetic cue was ignored. Ignoring the magnetic cue is a well-known phenomenon in magnetic research. For example, in a previous study, pigeons who underwent a clock-shift experiment flew only with deflection affected by the clock-shift and ignored the magnetic cue, regardless of the presence of bearing magnets (Ioalè et al., 2006).

Since the birds were prevented from using other navigational cues, such as celestial cues, during the current experiment, our assumption was that the only navigational cue available to them was the magnetic field (Bojarinova et al., 2023; Lefeldt et al., 2015). Furthermore, creating an induced magnetic field helps to isolate the tested individuals’ use of magnetoreception from their use of other signals. Unlike the results of this study, similar experimental setups, which included manipulation of the magnetic field using magnetic coils, have shown that birds left with only the use of the magnetic cue showed significant orientation towards the appropriate seasonal direction when no electromagnetic noise was present (Bojarinova et al., 2023; Zapka et al., 2009).

Taken together with the merged results of all three seasons, the significant within-individual directionality and the lack of significant orientation in the Control group lead us to speculate that there was some degree of electromagnetic noise in the induced magnetic field. The noise was indeed detected after recording the magnetic field continuously, rather than using point/average measurements, as we did to calculate the inclination angle. Although the power suppliers activated the coils properly and created the desired magnetic field, they eventually turned out to be unstable, leading to very slight fluctuations in the induced field that did not exist in the natural magnetic field. Individuals from the Control group may have detected these magnetic fluctuations and chosen to “ignore” their magnetic sense.

Recently, Bojarinova et al. (2023) showed that the avian magnetic compass has a sensitivity threshold to the oscillating magnetic field which is species-specific. For instance, the pied flycatcher (*Ficedula hypoleuca*) showed relatively low sensitivity, as individuals lost their magnetic orientation in an oscillating magnetic field with an amplitude of 190 nT and frequency of 1.41 MHz. In contrast to the pied flycatcher, the garden warbler (*Sylvia borin*) showed higher sensitivity, losing magnetic orientation in an oscillating magnetic field with an amplitude of 1–3 nT with the same frequency. It is plausible that the reed warbler may have higher sensitivity to oscillating magnetic fields, since this species is phylogenetically closer to the garden warbler than to the pied flycatcher.

When examining the results of the Treatment group we must take into account the possible sensitivity of the reed warblers to electromagnetic noise, and the negative effect of antibiotics on migratory restlessness orientation (Werber et al., 2022). In contrast to Werber et al. (2022) who exposed reed warblers to 48 h of antibiotics, in the current experiment the birds received a very short (12 h) antibiotic treatment. Apparently, this short treatment did not impair the magnetic sense completely. Treating birds with only two doses of antibiotics (and for a short duration) did not result in magnetic disorientation, but it seems that it reduced the birds’ sensitivity to electromagnetic noise. This could explain the relatively smaller directional variance in the Treatment group compared to the Control group. This could also explain the significant directionality of the Treatment group when merging the two spring seasons or all three experimental seasons. This is, of course, in contrast to the repeated lack of orientation in the Control group.

When using an unplugged coil (spring 2022), the birds from both groups showed significant southwest directionality. This is similar to the directionality seen in our Treatment group and to the Control group in a previous study conducted in our lab (Werber et al. 2022). Werber et al. (2022) used different experimental housing, located hundreds of meters away from the current experimental rooms. However, the individuals in the unplugged coil in our experiment and in the experiments conducted by Werber et al. (2022) were not subject to an induced (and fluctuating) magnetic field, and still showed significant southwest directionality. This supports the hypothesis that reduced sensitivity to electromagnetic noise led to increased directionality in the Treatment group.

The surprising southwest directionality, regardless of season, has repeated itself in four migration seasons and two separate studies. Unlike Werber et al. (2022), in the current experiment we manipulated the magnetic field, thus the southwest directionality was magnetic southwest and not geographic southwest. This supports the results of Werber et al. (2022) being linked to magnetic orientation and also prompts us to ask why this directionality is not season-dependent.

The results of this experiment are counterintuitive and differ from our initial expectations. However, in retrospect, examining the sensitivity of the birds to the electromagnetic noise could be a very sensitive and robust test of the magnetic sense (see also Bojarinova et al. 2023). Disorientation of the Control group could be explained by the sensitivity of the birds to electromagnetic noise. The seemingly “reverse” “positive”-effect of the short-term antibiotic treatment leads us to speculate that the antibiotics had an effect on magnetic-sense sensitivity but did not eliminate the sense completely.

The most obvious criticism on behavioral experiments that use antibiotics is the justified concern about unknown off-target effects (Bongers et al., 2022). This is also a reasonable concern for the current experiment. For instance, if the antibiotic treatment had resulted in a lack of jumps, leading to a lack of significant orientation, it would have been easy for us to attribute this behavior to a physiological side-effect of the antibiotic treatment. This is not the case in our experiment; on the contrary, considering the unexpected “positive” effect of short-term antibiotic treatment on orientation, attributing this behavior to an unknown side-effect seems less logical. Furthermore, previous results demonstrated that 48 hours’ antibiotic treatment led to an increase in directional variance in the Treatment group while having no effect on other behavioral and physiological measures such as feces production and activity levels (Werber et al., 2022).

Unlike Werber et al. (2022), we used a short-term treatment in the current experiment and we found an effect only on sensitivity, rather than on orientation. However, both results seem to share an effect of antibiotics on magnetic orientation, which changes with dosage. Still, in the current experiment, we cannot rule out an unknown side-effect of antibiotics contributing to our results. Furthermore, our interpretation of the results is an interpretation; other interpretations could be considered. However, considering the recent findings demonstrating the presence of MTB in animals (Natan et al., 2020), and the symbiotic magnetic sensing of protists using symbiotic MTB (Monteil et al., 2019), we believe that our experiment provides further support for the symbiotic magnetic sensing hypothesis (Natan et al., 2020; Natan & Vortman, 2017), but we are aware that this interpretation could be challenged. Moreover, the reported negative results (lack of significant orientation in the control group placed in a manipulated magnetic field) may motivate lack of publication. However such scientific approach leads to publication bias in Emlen funnel experiments. The best way to further examine the validity of our results, particularly since they are somewhat counterintuitive, is to reproduce such experiments in different labs.

Reproducibility is one of the underpinning principles of science. However, there is increasing concern about the reproducibility of studies in many fields (Laraway et al., 2019), specifically in the field of magnetoreception (Nimpf & Keays, 2022; Winklhofer et al., 2023). Different labs could easily adopt our experimental approach and examine the effect of various antibiotic dosages on magnetic sensitivity and reception. Eventually, unraveling a sense requires expertise in a range of disciplines. If indeed the underlying mechanism is symbiotic magnetic bacteria (Natan et al., 2020; Natan & Vortman, 2017), a multidisciplinary approach could unravel the mechanism behind this mysterious sense.

## Data availability

Upon acceptance all data will be available in Dryad data repository.

## References

1. Bojarinova, J., Kavokin, K., Cherbunin, R., Sannikov, D., Fedorishcheva, A., Pakhomov, A., & Chernetsov, N. (2023). Sensitivity threshold of avian magnetic compass to oscillating magnetic field is species-specific. Behavioral Ecology and Sociobiology, 77(1), 1–9. 10.1007/s00265-022-03282-7

2. Bongers, K. S., McDonald, R. A., Winner, K. M., Falkowski, N. R., Brown, C. A., Baker, J. M., Hinkle, K. J., Fergle, D. J., & Dickson, R. P. (2022). Antibiotics cause metabolic changes in mice primarily through microbiome modulation rather than behavioral changes. PLoS ONE, 17(3 March), 1–17. 10.1371/journal.pone.0265023

3. Chernetsov, N., Pakhomov, A., Kobylkov, D., Kishkinev, D., Holland, R. A., & Mouritsen, H. (2017). Migratory Eurasian Reed Warblers Can Use Magnetic Declination to Solve the Longitude Problem. Current Biology, 27(17), 2647–2651.e2. 10.1016/j.cub.2017.07.024

4. Emlen, S.T., Wiltschko, W., Demong, N.J., Wiltschko, R., & Bergman, S. (1976). Magnetic Direction Finding : Evidence for Its Use in Migratory Indigo Buntings. Science. 193(4252), 505–508. 10.1126/science.193.4252.505

5. Emlen, S. T., & Emlen, J. T. (1966). A Technique for Recording Migratory Orientation of Captive Birds. The Auk, 83(3), 361–367. 10.2307/4083048

6. Goswami, P., He, K., Li, J., Pan, Y., Roberts, A. P., & Lin, W. (2022). Magnetotactic bacteria and magnetofossils: ecology, evolution and environmental implications. Npj Biofilms and Microbiomes, 8(1), 1–14. 10.1038/s41522-022-00304-0

7. Gould, J. L. (1998). Sensory bases of navigation. Current Biology, 8(20), 731–738. 10.1016/s0960-9822(98)70461-0

8. Hore, P. J., & Mouritsen, H. (2016). The Radical-Pair Mechanism of Magnetoreception. Annual Review of Biophysics, 45, 299–344. 10.1146/annurev-biophys-032116-094545

9. Ioalè, P., Odetti, F., & Gagliardo, A. (2006). Do bearing magnets affect the extent of deflection in clock-shifted homing pigeons? Behavioral Ecology and Sociobiology, 60(4), 516–521. 10.1007/s00265-006-0194-0

10. Johnsen, S., Lohmann, K. J., & Warrant, E. J. (2020). Animal navigation: a noisy magnetic sense? Journal of Experimental Biology, 223(18), 1–7. 10.1242/JEB.164921

11. Kirschvink, J. L., Walker, M. M., & Diebel, C. E. (2001). Magnetite-based magnetoreception. Current Opinion in Neurobiology, 11(4), 462–467. 10.1016/S0959-4388(00)00235-X

12. Landler, L., Ruxton, G. D., & Malkemper, E. P. (2021). Advice on comparing two independent samples of circular data in biology. Scientific Reports, 11(1), 1–10. 10.1038/s41598-021-99299-5

13. Laraway, S., Snycerski, S., Pradhan, S., & Huitema, B. E. (2019). An Overview of Scientific Reproducibility: Consideration of Relevant Issues for Behavior Science/Analysis. Perspectives on Behavior Science, 42(1), 33–57. 10.1007/s40614-019-00193-3

14. Lefeldt, N., Dreyer, D., Schneider, N. L., Steenken, F., & Mouritsen, H. (2015). Migratory blackcaps tested in Emlen funnels can orient at 85 degrees but not at 88 degrees magnetic inclination. Journal of Experimental Biology, 218(2), 206–211. 10.1242/jeb.107235

15. Lefevre, C. T., & Bazylinski, D. A. (2013). Ecology, Diversity, and Evolution of Magnetotactic Bacteria. Microbiology and Molecular Biology Reviews, 77(3), 497–526. 10.1128/mmbr.00021-13

16. Lohmann, & Lohmann. (1994). Detection of Magnetic Inclination Angle By Sea Turtles: a Possible Mechanism for Determining Latitude. Journal of Experimental Biology, 194(1), 23–32. 10.1242/jeb.194.1.23

17. Mathis, A., Mamidanna, P., Cury, K. M., Abe, T., Murthy, V. N., Mathis, M. W., & Bethge, M. (2018). DeepLabCut: markerless pose estimation of user-defined body parts with deep learning. Nature Neuroscience, 21(9), 1281–1289. 10.1038/s41593-018-0209-y

18. Meyer, F., Paarmann, D., D’Souza, M., Olson, R., Glass, E. M., Kubal, M., Paczian, T., Rodriguez, A., Stevens, R., Wilke, A., Wilkening, J., & Edwards, R. A. (2008). The metagenomics RAST server - A public resource for the automatic phylogenetic and functional analysis of metagenomes. BMC Bioinformatics, 9, 1–8. 10.1186/1471-2105-9-386

19. Monteil, C. L., Vallenet, D., Menguy, N., Benzerara, K., Barbe, V., Fouteau, S., Cruaud, C., Floriani, M., Viollier, E., Adryanczyk, G., Leonhardt, N., Faivre, D., Pignol, D., López-García, P., Weld, R. J., & Lefevre, C. T. (2019). Ectosymbiotic bacteria at the origin of magnetoreception in a marine protist. Nature Microbiology, 4(7), 1088–1095. 10.1038/s41564-019-0432-7

20. Natan, E., Fitak, R. R., Werber, Y., & Vortman, Y. (2020). Symbiotic magnetic sensing: raising evidence and beyond. Philosophical Transactions of the Royal Society of London. Series B, Biological Sciences, 375(1808), 20190595. 10.1098/rstb.2019.0595

21. Natan, E., & Vortman, Y. (2017). The symbiotic magnetic-sensing hypothesis: Do Magnetotactic Bacteria underlie the magnetic sensing capability of animals? Movement Ecology, 5(1). 10.1186/s40462-017-0113-1

22. Nath, T., Mathis, A., Chen, A. C., Patel, A., Bethge, M., & Mathis, M. W. (2019). Using DeepLabCut for 3D markerless pose estimation across species and behaviors. Nature protocols, 14(7), 2152–2176.

23. Nimpf, S., & Keays, D. A. (2022). Myths in magnetosensation. IScience, 25(6), 104454. 10.1016/j.isci.2022.104454

24. Nordmann, G. C., Hochstoeger, T., & Keays, D. A. (2017). Unsolved mysteries: Magnetoreception—A sense without a receptor. PLoS Biology, 15(10), 1–10. 10.1371/journal.pbio.2003234

25. Packmor, F., Kishkinev, D., Bittermann, F., Kofler, B., Machowetz, C., Zechmeister, T., Zawadzki, L. C., Guilford, T., & Holland, R. A. (2021). A magnet attached to the forehead disrupts magnetic compass orientation in a migratory songbird. Journal of Experimental Biology, 224(22). 10.1242/jeb.243337

26. Pearson, D. J., Kennerley, P. R., & Small, B. J. (2002). Eurasian Reed Warbler: the characters and variation associated with the Asian form fuscus. British Birds, 95:42–61

27. Putman, N. F., Endres, C. S., Lohmann, C. M. F., & Lohmann, K. J. (2011). Longitude perception and bicoordinate magnetic maps in sea turtles. Current Biology, 21(6), 463–466. 10.1016/j.cub.2011.01.057

28. Schwarze, S., Schneider, N. L., Reichl, T., Dreyer, D., Lefeldt, N., Engels, S., Baker, N., Hore, P. J., & Mouritsen, H. (2016). Weak broadband electromagnetic fields are more disruptive to magnetic compass orientation in a night-migratory songbird (Erithacus rubecula) than strong narrow-band fields. Frontiers in Behavioral Neuroscience, 10(MAR), 1–13. 10.3389/fnbeh.2016.00055

29. Wang, C. X., Hilburn, I. A., Wu, D.-A., Mizuhara, Y., Cousté, C. P., Abrahams, J. N. H., Bernstein, S. E., Matani, A., Shimojo, S., & Kirschvink, J. L. (2019). Transduction of the Geomagnetic Field as Evidenced from Alpha-band Activity in the Human Brain. ENeuro. 10.1523/ENEURO.0483-18.2019

30. Werber, Y., Natan, E., Lavner, Y., & Vortman, Y. (2022). Antibiotics affect migratory restlessness orientation. Journal of Ethology, 40(2), 175–180. 10.1007/s10164-022-00747-0

31. Wiltschko, W., & Wiltschko, R. (1972). Magnetic Compass of European Robins. Science, 176, 62–64.

32. Wiltschko, W., & Wiltschko, R. (2012). Global navigation in migratory birds: Tracks, strategies, and interactions between mechanisms. Current Opinion in Neurobiology, 22(2), 328–335. 10.1016/j.conb.2011.12.012

33. Winklhofer, M., Hore, P. J., & Mouritsen, H. (2023). No evidence for magnetic field effects on the behaviour of Drosophila. *March*. 10.1038/s41586-023-06397-7

34. Wynn, J., Padget, O., Mouritsen, H., Morford, J., Jaggers, P., & Guilford, T. (2022). Magnetic stop signs signal a European songbird&#x2019;s arrival at the breeding site after migration. Science, 375(6579), 446–449. 10.1126/science.abj4210

35. Zapka, M., Heyers, D., Hein, C. M., Engels, S., Schneider, N. L., Hans, J., Weiler, S., Dreyer, D., Kishkinev, D., Wild, J. M., & Mouritsen, H. (2009). Visual but not trigeminal mediation of magnetic compass information in a migratory bird. Nature, 461(7268), 1274–1277. 10.1038/nature08528

