## supplementary material for "Short-term antibiotics, induced magnetic field, and magnetic sensitivity in a migratory songbird"

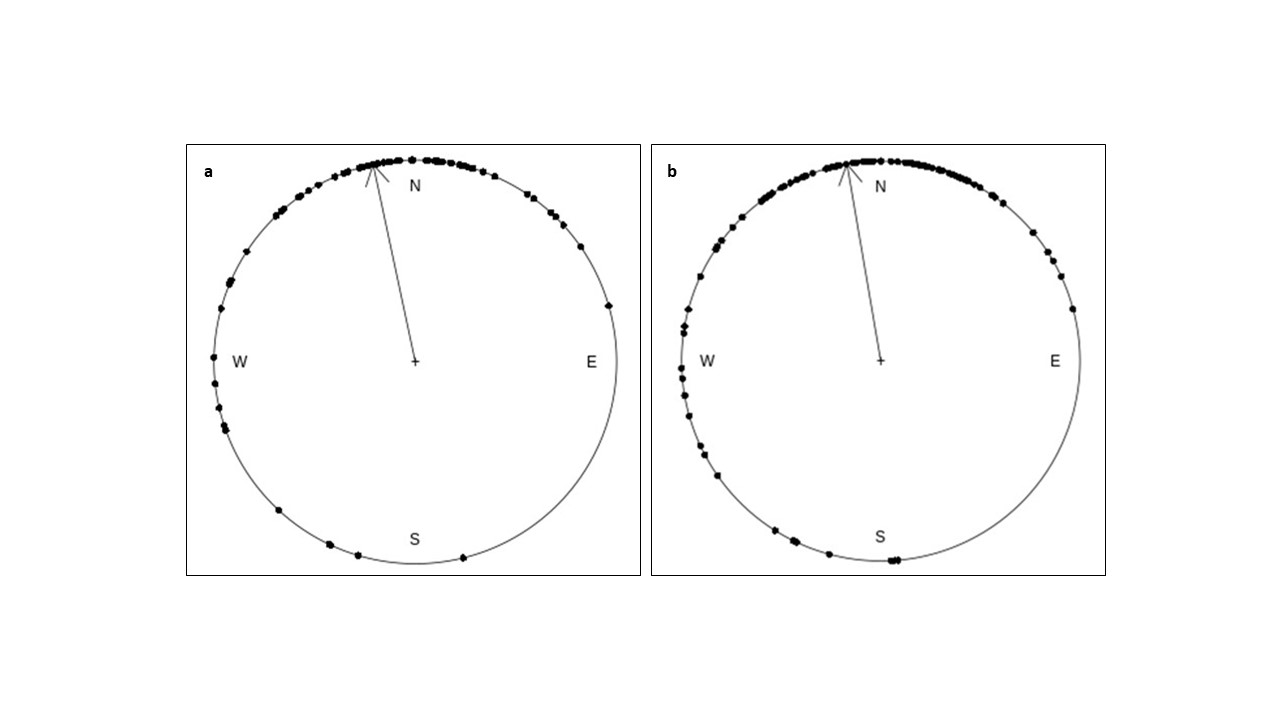

Fig. S1 Comparison between orientation graphs of an individual (ring number - X552113), using data obtained from Deep Lab Cut video analysis (Azimuth – 347°, R- 0.68, p<0.0001; a), and manual video analysis (Azimuth – 350°, R- 0.723, p<0.0001; b).
